# Comparison of starch granule sizes of maize seeds between parental lines and derived hybrids suggests a maternal inheritance trend

**DOI:** 10.1101/2021.02.03.429493

**Authors:** Liangjie Niu, Wei Wang

## Abstract

Maize (*Zea mays*) starch is an important agricultural commodity that serves as food, feed, and a raw material for industrial purposes. It is organized into starch granules (SG) inside amyloplasts and is highly accumulated in endosperms. Maize hybrids, which exhibits heterosis over their parents, are globally grown due to higher vigor of the F1 plants. However, the parental effect on the size of SG in F1 hybrid seeds remains unclear. Here we compared the seed SG sizes among two parental inbred lines (Chang7-2 and Zheng58) as well as their reciprocal hybrids. SG was observed *in situ* and *in vitro* with SEM. The size of seed SG in hybrids was more like that of female parents, especially for large SG population. Thus, the control of SG size exhibits a maternal inheritance trend in the context of plastid (amyloplast) inheritance. Our results provide some insight on selecting parental inbred lines for breeding maize hybrids with different SG sizes.

## INTRODUCTION

Starch is the predominant carbohydrate reserve in plants (Zeeman et al., 2010). It consists of glucose residues linked to each other by α-1,4-linkages (amylose) with occasional α-1,6-branches (amylopectin) (Zeeman et al., 2010). In addition to as food and feed, starch is used as texturizer, gelling agent, thickener, adhesive, moisture-retainer, biofuels etc. (Niu et al., 2019). Starch accumulates in amyloplasts in a semi-crystalline granular form, which is divided into two forms: storage starch and transient starch (Zeeman et al., 2010). Storage starch granule (SG) mainly exists in cereal seeds, storage roots, and tubers, with a highly varied size of 2 to 50 μm dependent on plant species (Seung and Smith, 2019). Transient SG is accumulated in leaf chloroplast and is usually small (∼2 μm) with a uniformly flattened disc shape among plant species (Seung et al., 2017; Yu et al., 2017; Zeeman et al., 2002). The SG with different sizes can be used for specific applications. Large SG with more amylose, higher peak, trough and final pasting viscosities, and lower gelatinization temperature than small SG (Cai et al., 2014), resulting in different bread- and noodle-making quality (Zi et al., 2019). Fine SG can be used for generating thin films (Lindeboom et al. 2004) and fatty mouthfeel (Ma et al., 2006).

Many studies have demonstrated that the morphology and size of storage SG are mainly determined by species origin (Zeeman et al., 2010), biochemistry and physiology of plants (Gregorová et al., 2006; Alcázar-Alay and Meireles., 2015; Zhao et al., 2018). Some QTLs and protein (e.g. SUBSTANDARD STARCH GRAIN6, starch synthase I) controls SG size have been identified in maize kernels, rice and wheat endosperm using mutant (Liu N et al. 2018; Matsushima R et al. 2016;McMaugh SJ et al. 2014). However, little reports about the effect of parental inbred lines on the seed SG size in derived hybrids in maize.

Maize (*Zea mays*) is globally grown for its high seed starch content. Its hybrids exhibit heterosis over parental lines in plant growth and development, yield and stress tolerance (Hoecker et al., 2006). The hybrid kernel traits, especially weight and size, are mostly affected by female parents (Zhang et al., 2016). Here two inbred lines Chang7-2 (C7-2) and Zheng58 (Z58) and their derived hybrids were used to instigate the parental effect on the SG sizes by scanning electron microscopy.

## MATERIALS AND METHODS

Maize inbred lines C7-2 and Z58 were grown and reciprocally crossed as described previously (Ning et al., 2017). Mature seeds of C7-2, Z58, C7-2 × Z58, and Z58 ×C7-2 were used here. Scanning electron microscopy (SEM) observation of seed SG *in situ* and *in vitro* was exactly performed as our recent work (Niu et al. 2019). The SG size was expressed as spherical diameter by measuring at least 300 granules. All assays were conducted in at least three biological replicates. Statistically significant differences were accessed by the Student’s t-test (p<0.05).

## RESULTS

The seeds of C7-2 and Z58 significantly differ in shape, size and weight. The C7-2 seeds were like triangle and longer, whereas the Z58 seeds were rounder and wider (Fig. 1a-1c). The derived hybrids were like the female parent in morphology but with an increased size and weight (Fig. 1d-1e). Moreover, the proportion of the endosperm in the hybrid seeds was determined by the female parent (Fig. 1f).

**Figure 1.**
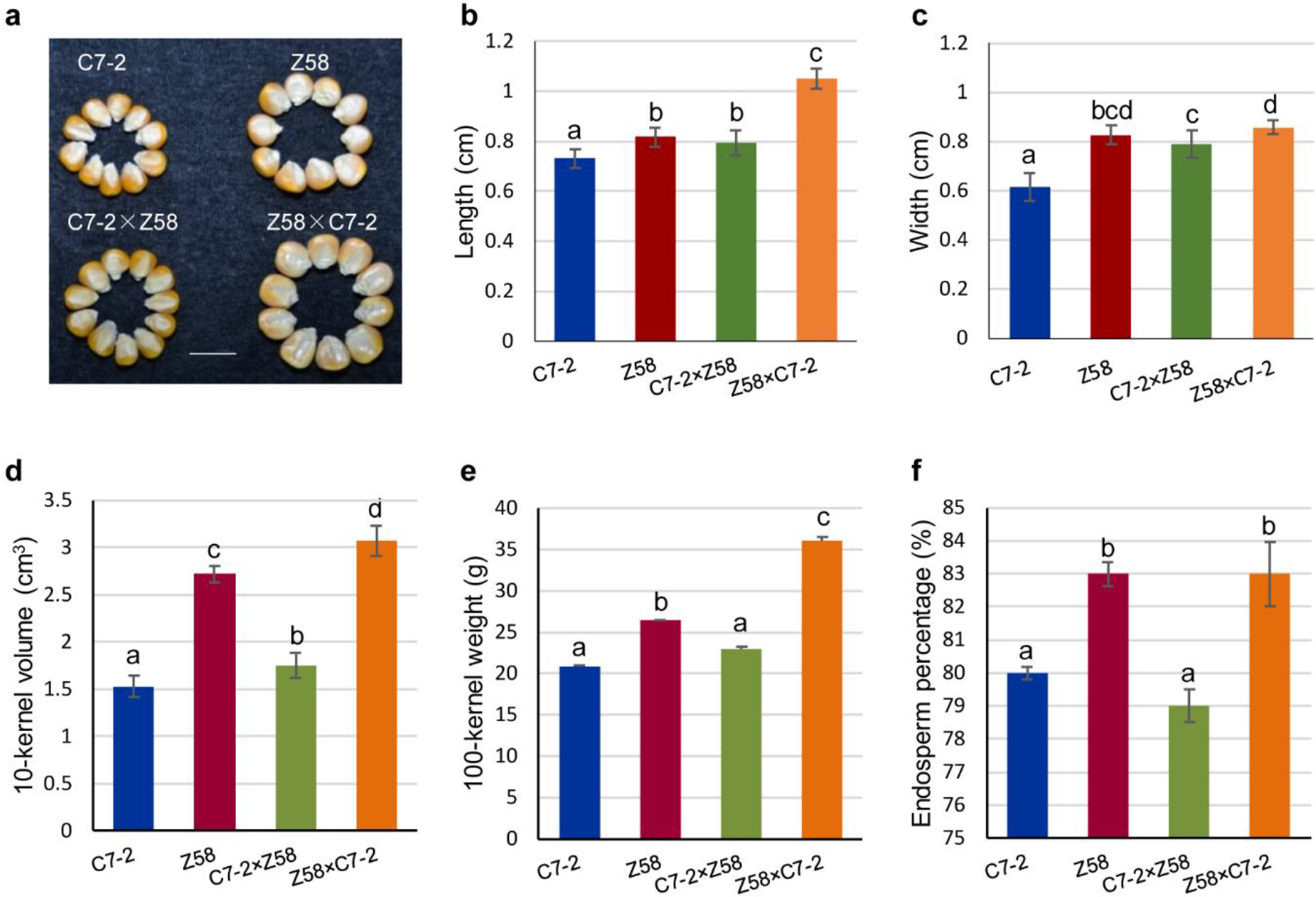
Phenotypic analysis of seeds. **a** Seeds of two inbred lines and their reciprocal hybrids. Bar = 1 cm. **b** The length of kernels (mean ±SD, n=50). **c** The width of kernels (mean ±SD, n=50). **d** The 10-kernel volume (mean ±SD, n=5). **e** The 100-grain weight (mean ±SD, n=5). **f** The ratio of endosperm in seed (mean ±SD, n=5).

The seed SG *in situ* of the four materials were in spherical shape (Fig. 2a), but the SG surface of the parental seeds was smooth but was rough in the hybrids with more membrane debris. The size of SG in all samples ranged from 4.5 to 19.5 μm, and the hybrid seeds were more similar to the female parent in the average SG size and distribution (Fig. 2b-2c), especially the ratio of large SG population (Fig. 3d), exhibiting a maternal inheritance trend.

**Figure 2.**
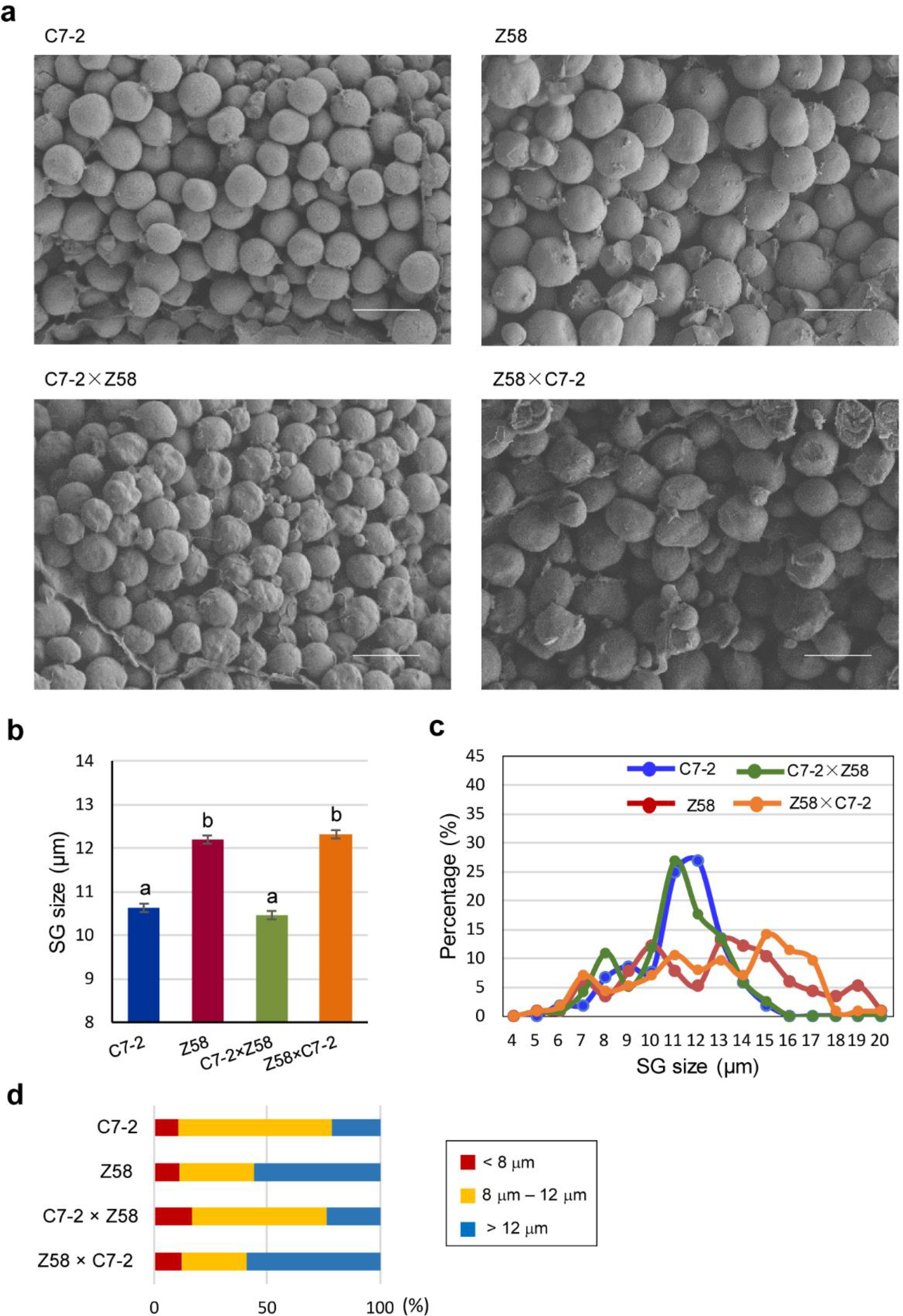
The morphology and size of seed SG *in situ*. **a** SEM observation. Bar = 20 μm. **b** Average SG diameter of. **c** SG size distribution. **d** The size change in different SG populations.

**Figure 3.**
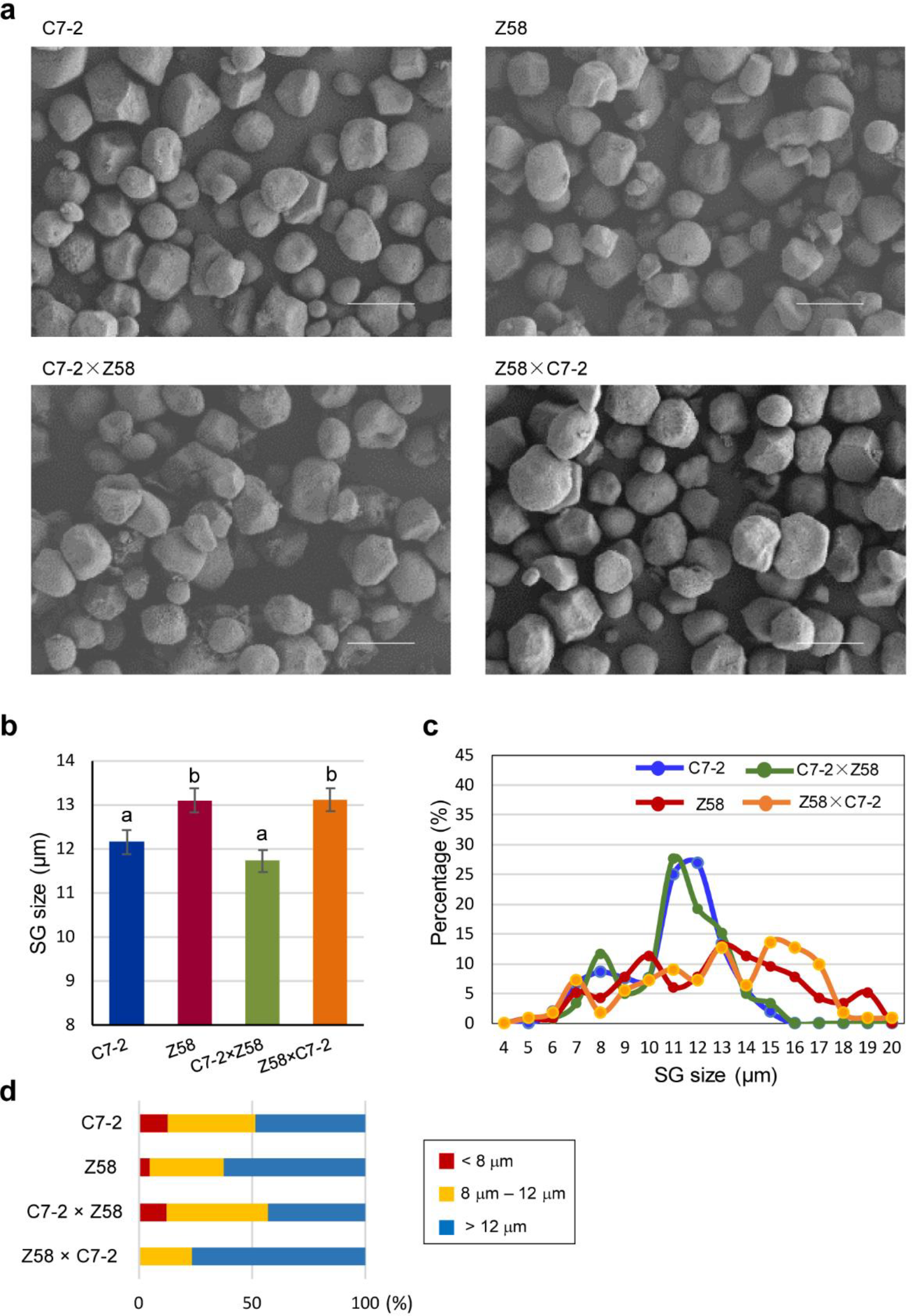
The morphology and size of seed SG in vitro. **a** SEM observation. Bar = 20 μm. **b** Average SG diameter of. **c** SG size distribution. **d** The size change in different SG.

The seed SG *in vitro* in all samples were mostly polygonal with similar surface features, because the membrane debris on the surface of the SG were removed during the isolation (Fig. 3a). The size of the isolated SG varied from 4.7 to 27.98 μm (Fig. 3b), greater than that *in situ*. Even so, the size distribution of the SG *in vitro* was comparable to the *in vivo* results (Fig. 3d). Additionally, the SG size correlated with the 100-kernel weight and the endosperm sizes, with correlation coefficients of 0.5595 (p<0.05) and 0.5773 (p<0.05), respectively.

## DISCUSSION

Maize seeds consist of up to 78% starch that is overwhelmingly accumulated in the endosperm. The endosperm plays a key role in determining seed size (Berger et al., 2006). Large SG is the major determinant of wheat endosperm weight (Chojkecki et al., 1986). Female parent significantly influences maize hybrid seed traits through altering endosperm cell number (Jones et al., 1996).

## DISCUSSION

Maize seeds consist of up to 78% starch that is overwhelmingly accumulated in the endosperm. The endosperm plays a key role in determining seed size (Berger et al., 2006). Large SG is the major determinant of wheat endosperm weight (Chojkecki et al., 1986). Female parent significantly influences maize hybrid seed traits through altering endosperm cell number (Jones et al., 1996).

In the present study we found the SG size in hybrid seeds was more affect by the female parent. Two reasons may explain this phenomenon. First, the endosperm is a triploid tissue that receives two genomes from mother and one genome from the paternal side (Li and Li, 2015). Gómez et al. (2002) found endosperm transfer cell can facilitate the transport of maternal solutes and nutrients at the interface between maternal tissues and the endosperm. Second, starch is synthesized and stored in plastids (chloroplasts/amyloplasts) in leaves or seeds (Seung and Smith, 2019). Plastids are maternally inherited in plants, as observed in Epilobium plants by electron microscopy (Schmitz and Kowallik 1987). It is suggested that maternal plastid inheritance is the elimination of paternal plastids during fertilization or the exclusion of plastids from germ cells during pollen development (Kuroiwa 2010). However, the mechanisms of parental inbred lines on the SG size in their hybrid seeds remains unclear in the case of maize. The size and shape of the SG are controlled by multiple factors, such as botanical species origin, biochemistry and physiology of plants, and SG properties (Ellis et al., 1998; Nordmark and Ziegler., 2002; Ji et al., 2003a, 2003b; Lu et al., 2011; Cai et al., 2014). Super-sweet, pop, waxy, and dent maize differ significantly in size and shape of seed SG (Cui et al., 2014). The SG size and morphology in maize seeds are affected by nitrogen levels (Zhao et al., 2018). A significant SNPs within genes GRMZM2G419655 and GRMZM2G511067 was identified as functional markers in screening maize SG size (Liu et al., 2018). In rice, amyloplast membrane protein SUBSTANDARD STARCH GRAIN6 can control SG size in endosperm (Matsushima et al., 2016). In wheat, starch synthase I can change the morphology, size and fine structure of starch in wheat endosperm by affecting the amylose content (McMaugh et al., 2014). However, the mechanism controlling the size and shape is largely unclear.

It is needed to note that maize seed SG were generally spherical *in situ* with and polygonal *in vitro* with comparative sizes as observed previously (Li et al. 2007; Zhang et al., 2011; Paraginski et al., 2014; Zhao et al., 2018; Niu et al., 2019). The large SG took a slight greater proportion in the isolated SG, possibly because some of the fine SG was lost during the isolation and all SG may somewhat swell from water-absorbing during the isolation. In addition, only two elite inbred lines C7-2 and Z58 in China were used in the present study and their SG size was different but comparable, if the SG size of the parental lines were greatly varied, the derived hybrids may varied greatly in seed SG sizes.

We found that the size of seed SG in hybrids was more like that of female parents, especially for large SG population, exhibiting a maternal inheritance trend. This provides some insight on selecting parental inbred lines for breeding maize hybrids with different SG sizes for different applications.

## Abbreviations

SG: starch granules

